# How to get the most bang for your buck: the evolution and physiology of nutrition-dependent resource allocation strategies

**DOI:** 10.1101/113027

**Authors:** Enoch Ng’oma, Anna M. Perinchery, Elizabeth G. King

## Abstract

All organisms utilize resources to grow, survive, and reproduce. The supply of these resources varies widely across landscapes and time, imposing ultimate constraints on the maximal trait values for allocation-related traits. In this review, we address three key questions fundamental to our understanding of the evolution of allocation strategies and their underlying mechanisms. First, we ask: how diverse are flexible resource allocation strategies among different organisms? We find there are many, varied, examples of flexible strategies that depend on nutrition. However, this diversity is often ignored in some of the best-known cases of resource allocation shifts, such as the commonly observed pattern of lifespan extension under nutrient limitation. A greater appreciation of the wide variety of flexible allocation strategies leads directly to our second major question: what conditions select for different plastic allocation strategies? Here, we highlight the need for additional models that explicitly consider the evolution of phenotypically plastic allocation strategies and empirical tests of the predictions of those models in natural populations. Finally, we consider the question: what are the underlying mechanisms determining resource allocation strategies? Although evolutionary biologists assume differential allocation of resources is a major factor limiting trait evolution, few proximate mechanisms are known that specifically support the model. We argue that an integrated framework can reconcile evolutionary models with proximate mechanisms that appear at first glance to be in conflict with these models. Overall, we encourage future studies to 1) mimic ecological conditions in which those patterns evolve, and 2) take advantage of the ‘omic’ opportunities to produce multi-level data and analytical models that effectively integrate across physiological and evolutionary theory.

## 1 The central importance of the interplay between resource acquisition and allocation

The amount of resources available to organisms, whether the source is sunlight, plant matter, or prey animals, is inherently variable over the landscape and across time. This variability presents a fundamental challenge to all organisms, from the smallest microorganisms to the largest plants and animals, all of which must coordinate the acquisition of resources from the environment with allocation of those resources among the many competing functions and structures that contribute to the organisms’ fitness. When faced with variation in available resources, individuals could respond in one of two ways: (1) maintaining the same relative proportion allocated to each trait or (2) exhibiting phenotypic plasticity in resource allocation by altering the relative amount of resources allocated to one trait versus others. When the optimal allocation strategy changes with resource availability, selection will favour the evolution of a phenotypically plastic allocation strategy.

The inescapable link between the amount of resources available to an organism and subsequent allocation of those resources means it is critical to consider how allocation strategies change across a range of resource availabilities. There are many examples of flexible strategies that depend on availability. For example, an adaptive shift in resource allocation is thought to underlie the commonly observed pattern of lifespan extension under dietary restriction (reviewed in [1–5]. Likewise, sexually selected traits often show strong condition dependence (i.e. dependence on acquisition), also thought to result from an adaptive shift in allocation (reviewed in [6,7]. Even the current obesity epidemic in modern human populations is often hypothesized to result from a mismatch between a selective environment favouring increased storage under high resources and the modern environment of constant high resource availability [8] (see [9] for a recent review). To understand this wide diversity in allocation strategies in the natural world, we must understand how different ecological conditions select for different strategies and what mechanistic changes underlie these strategies.

Understanding how and why this coordination of resource allocation with availability evolves has implications for nearly all areas of biology. Energetic costs to biological structures and functions (i.e. allocation trade-offs) are assumed to be universal and a major factor limiting trait evolution [10,11]. Typically, less attention is focused on the role of variation in the acquisition of resources, though it is no less important in determining trait values, and can obscure the detection of functional trade-offs. In a seminal paper, van Noordwijk and de Jong introduced the Y model - a mathematical model linking resource acquisition and resource allocation [12], which has been a central concept in the field of life history evolution. In the Y model, two traits draw from a single resource pool, with trait values determined by the proportion of resources allocated to each (Figure 1 [12]). One of the key strengths of this model is its simplicity and generality; it can be applied to diverse questions such as why and how organisms age, what limits crop yields in different environments, why some species produce hundreds of offspring while others produce very few, and what constrains the evolution of fitness. While the Y model provides a conceptual starting point to understand the evolution of acquisition and allocation, in the Y model the underlying mechanisms governing these processes are treated as a black box. Likewise, our empirical knowledge of the genetic and physiological mechanisms underlying these processes is still limited, due in large part to their vast complexity [13,14]. The allocation of resources is thought to influence nearly all the major structures and functions of an organism, is affected by an array of interacting physiological pathways, is variable across the lifetime of the organism, and interacts with many different environmental factors. To achieve a complete understanding of how resource allocation trade-offs govern these processes, we must explicitly consider its interaction with resource acquisition and integrate across genomics, physiology, and evolution.

**Figure 1:**
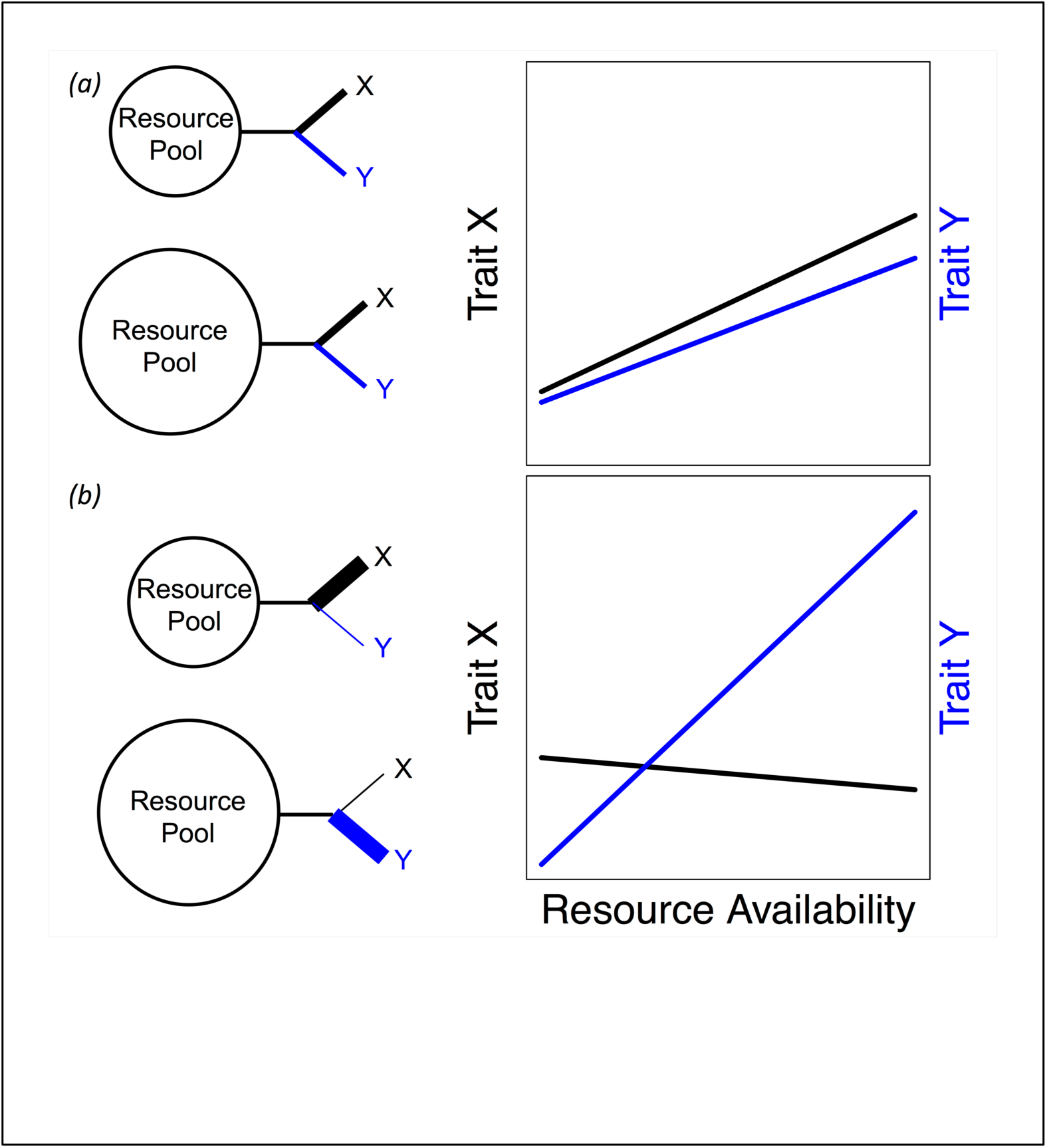
Expectations for trait values for two traits involved in a resource allocation trade-off when A) there is no phenotypic plasticity in allocation in response to resource availability, and B) there is phenotypic plasticity in allocation with increasing proportions allocated to trait Y as resource availability increases.

As we advance our ability to collect “omic” data at multiple levels (genomics, transcriptomics, proteomics, metabolomics, etc.) and in multiple environments, achieving this integration is becoming increasingly feasible. A major challenge now is developing new analytical methods to address multi-level, multi-environment data, and pulling out emergent themes that will help us better understand the complex processes underlying trade-offs and linking these with evolutionary models. We argue that resource allocation is a natural focal point in this effort. This relatively straightforward concept has the potential to integrate knowledge across fields and address key questions facing the intersection between evolutionary and molecular biology.

In this review, our goals are to: 1) detail the diversity of resource allocation strategies in response to environmental fluctuations in resource availability, 2) review the evolutionary explanations for these strategies and highlight where new models are needed, and 3) evaluate current approaches and suggest strategies for understanding the genetic and physiological mechanisms underlying resource allocation strategies.

## 2 The diversity of phenotypically plastic resource allocation strategies in the natural world

In the wild, organisms vary widely between species and populations in how they respond to variation in resource acquisition, with a diverse array of examples of phenotypically plastic resource allocation strategies (Table S1). Variation in resource acquisition can result from variation in resource abundance in the environment, and/or from differences among individuals in their ability to acquire resources. By far the largest challenge in describing broad patterns of phenotypic plasticity in allocation strategies is to directly quantify resource acquisition and the amount of those resources allocated to different traits. In only a very few cases have resource acquisition and allocation been successfully estimated in terms of energy units (e.g. [15–18]). In the majority of studies, these patterns must instead be inferred indirectly from phenotypic patterns.

The problem of estimating acquisition can be avoided in part when acquisition can be experimentally manipulated via resource restriction. When resource levels are restricted, the expectation for resource-based trait values is that they will also decrease. When trait values increase instead or remain constant, it suggests increased allocation to that trait (Figure 1). A well-examined example of this type of pattern is the commonly observed increase in lifespan (hypothesized to be due to increased allocation to somatic maintenance) under food restriction coupled with reduced reproduction (reviewed in [1–5]). The majority of the work on the response of lifespan to food restriction has been focused on model organisms. While there are several examples in non-model species that show a similar response (Table S1), not all species live longer on food restriction [19], including some species of water striders [20], house flies [21], squirrel monkeys [22], and rotifers [23,24]. Additionally, several species show a marked *increase* in reproductive allocation under low resource conditions (flatworms [25], guppies [26], rotifers [24]), demonstrating reproductive allocation does not always decrease under food restriction. Another trade-off that is particularly well characterized in terms of differential resource allocation is the trade-off between flight capability and reproduction in several wing dimorphic insect species (reviewed in [27–30]). In these species, there exist discrete flight capable (macropterous) and flightless (micropterous or apterous) morphs. Wing morphology displays phenotypic plasticity in response to several environmental variables including rearing density, a likely correlate with acquisition, with different species displaying very different responses. In aphids and planthoppers, induction of flight capable morphs increases in response to crowding and low nutrition [31,32], while in crickets, group rearing and other stressors increase induction of flightless morphs [33,34]. Both of these examples, the lifespan-reproduction trade-off and the flight capability-reproduction trade-off, demonstrate the wide variation in allocation patterns across different species.

Most experimental manipulations of acquisition simply consider a single “low” and single “high” resource environment, and often the diet used is artificial and quite different from the organism’s natural diet. Recently, the community has begun to take a “nutritional geometry” perspective, considering wider ranges of nutritional conditions, both in terms of caloric content and individual diet components (i.e. protein, carbohydrate, and lipid content), as well as a wider range of the timing of resource level changes across an organism’s lifetime [35–37]. These efforts provide a much more complete picture of how an organism responds to diet, in that they distinguish between allocation changes due to limitation in specific nutrient classes vs effects due to more general caloric restriction [38,39]. However, this approach increases complexity, which can make interpreting the results in an evolutionary context a challenge when patterns are highly dynamic. To best place diet manipulations in an evolutionary context, we need ecological studies that characterize typical diet sources, and the degree of natural variation in resource availability experienced by populations in the wild. For many populations, this goal will be a challenge.

Overall, a broad view of trait variation reveals many examples of variation in plastic resource allocation in response to variation in acquisition (Table S1). Often, patterns vary substantially among closely related species (e.g. [20,22,23,25]), among populations of the same species (e.g. [20,26]), or between different inbred strains [40]. These examples argue against any hard and fast, universal resource allocation strategies in response to variation in acquisition and lead to the key questions of why and how environmental variation in resource availability leads to the evolution of different resource allocation strategies. From the resource allocation strategies detailed in Table S1, we can conclude two basic points: 1) phenotypic plasticity in resource allocation is common, and 2) the pattern of plasticity varies widely among populations and taxa. Beyond these points, it is difficult to draw any general conclusions given that the dataset is biased (e.g. model organisms are overrepresented), and there are many different methods for estimating resource allocation (see discussion of these methods above), making it challenging to generalize across studies. Clearly, we require a better understanding of the ecological conditions that would lead to the evolution of such different populations.

## 3 Why do phenotypically plastic resource allocation strategies evolve?

There is a long and rich history of theoretical evolutionary models addressing both optimal resource allocation patterns in different environmental conditions (i.e. life history evolution models; e.g. [41–45]; see [10,46] for extensive reviews), and the evolution of phenotypic plasticity [47–53]. However, there are few models that specifically focus on the evolution of phenotypically plastic resource allocation in response to variation in resource availability [54–56]. While this category might seem to be a special case, there is reason to expect general models of phenotypic plasticity might not be fully applicable to variation in resource availability. Resource availability places an ultimate constraint on the maximal trait values for allocation-related traits, and in that way, it is fundamentally different from other types of environmental conditions. The dependency creates the somewhat paradoxical situation in which no plasticity in allocation will lead to plasticity in trait values, as they will necessarily decrease with resource availability (Figure 1). Thus, it is critical for theoretical models to explicitly consider variability in resource availability when predicting how plastic allocation strategies will evolve.

One emergent property of models that do explicitly consider the interplay between acquisition and allocation is that environmental predictability (i.e. whether current resource availability is correlated with future availability) is a major determinant of the pattern of phenotypic plasticity that evolves [54–56]. In a model considering allocation to flight capability versus reproduction, King *et al* [54] showed completely opposite patterns of plasticity in allocation are expected to evolve in environments with predictable versus unpredictable patterns of resource availability. Fischer and co-workers [55,56] showed that, in response to short term resource availability fluctuations, populations should evolve to allocate toward somatic maintenance under low food conditions. However, this response is more complicated. If conditions are low enough to be indicative of low survival probability, allocation to survival is not favoured. Rather, a terminal investment strategy, investing heavily in reproduction at the expense of survival, is favoured.

One area where models of the evolution of condition-dependent (i.e. acquisition-dependent) resource allocation strategies is well developed is in the field of sexual signalling. In many cases, male advertisements to females are dependent on the condition of the male, producing so-called ‘honest’ signals (e.g. [57,58]; for reviews see [59–61]. This condition dependence can be continuous (e.g. call duration in male grey tree frogs [57]) or a discrete polymorphism (e.g. sexually dimorphic mandible growth in stag beetles [62]). There are several models considering how the benefits and costs of increased allocation toward a sexual signal change depending on an individual’s condition [59,61,63], with models predicting low condition individuals that allocate more toward sexual signals experience lower benefits and/or higher costs depending on the assumptions of the model (see [61]). These models are a subset of models considering allocation strategies in poor condition as a ‘best of a bad lot’ strategy [46]. In essence, it does not pay to invest heavily in a sexual signal if one simply does not have enough resources to produce a high-quality signal that will attract many mates.

The majority of evolutionary models focus solely on why, not how, allocation patterns evolve, ignoring the underlying mechanisms. Often, this is a sensible strategy, given that when mechanisms don’t act as ultimate constraints, evolutionary endpoint will remain the same, irrespective of the specifics of the mechanistic underpinning. Nevertheless, evolutionary models that incorporate explicit mechanisms, can be highly informative in explaining the mechanisms underlying evolutionary patterns. For example, Mangel and Munch [64] integrated physiological parameters such as oxidative damage associated with faster growth and resource allocation to damage repair in a model predicting when compensatory growth (increased allocation to growth following a period of food restriction) should evolve. Only by explicitly incorporating the physiological mechanisms of damage and repair, were they able to simulate patterns of compensatory growth that matched observations. Compensatory growth never arose using a simple optimality framework, demonstrating that explicitly incorporating physiology can fundamentally change the predictions of life history models in some cases. We encourage the development of evolutionary models that integrate proximate mechanisms as a way to expand our understanding of the evolution of resource allocation strategies in multiple systems.

## 4 Genetic and physiological mechanisms underlying phenotypic plasticity in resource allocation

It is clear organisms have evolved the ability to shift the allocation of resources in response to their nutritional state in many different ways, but *how* do they accomplish this change? What physiological changes accompany a shift in allocation strategy and what genes are involved? Not surprisingly, the greatest progress in the effort to uncover the mechanisms governing the coordination between acquisition and allocation comes from model organisms (e.g. yeast, worms, flies, and mice) that have been the focus of studies for decades. However, the relatively recent “omic” technologies available, and the decreasing cost of these technologies, make it increasingly feasible to gather data at multiple levels of the genotype to phenotype map in multiple environments for nearly any organism, opening up the possibility of moving beyond unnatural manipulations in model organisms and toward more ecologically relevant contexts.

### a Evolutionary endocrinology suggests key role of hormones in resource allocation

At first glance, resource acquisition and allocation might seem hopelessly complex, casting doubt on the prospect of uncovering the proximate mechanisms involved in the relatively subtle variation, at least when compared to mutants, in natural populations. However, an emergent theme from several systems, including many of the above detailed examples in model organisms, is the key role of hormone pathways as major determinants of resource allocation. These discoveries have spurred the expansion of the field of “evolutionary endocrinology” [65–68]. For instance, we have learned a great deal about the mechanisms governing allocation of resources in response to environmental changes from genetic screens and mutational analysis in model organisms (e.g. [68–70]). In this section, we review our current knowledge in model and non-model systems on how plasticity in nutrient allocation and hormonal signalling impact reproduction-lifespan and reproduction-dispersal trade-offs. The hormone pathways we discuss here include insulin, ecdysone, and juvenile hormone.

#### i Large effect mutations support the idea that major signalling pathways modulate allocation

Studies that have yielded the most insights have tended to focus on mutations of large effect. Several studies have implicated leptin as a mediator of acquisition and allocation of nutrients. In mammals, leptin together with AMPK (AMP-activated protein kinase) control appetite thus regulating nutrient intake [71], and leptin also mediates the energetic trade-off of reproduction with the immune system by acting as a proximate endocrine indicator of the energy state to the immune system [72]. In *D. melanogaster*, Upd2 (unpaired 2), a functional homolog of leptin [73], causes a nutrient dependent effect on growth, mediating production of Dilps (*Drosophila* insulin-like peptides) in the fed state, and subsequent secretion of insulin in response to dietary fat [73]. These studies demonstrate a direct connection between nutrient limitation and allocation.

Both ecdysone and the insulin/insulin-like signalling pathway (IIS) have a role in the plastic allocation of nutrients. Sequential perturbation of IIS and ecdysone signalling in ovarian somatic cells of *D. melanogaster* on different diets showed that ecdysone signalling regulated the rate of increase in ovary volume in general while IIS conferred the same effect before larvae attained critical weight [74,75]. This nutrient-dependent development of the ovary illustrates the role of hormonal signalling in plastic allocation of nutrients. Perhaps one of the most significant contributions emerging from mutation studies is that IIS signalling pathways are critical in the regulation of lifespan in many species. In several model organisms (including fly, mice and worm), reduced IIS phenocopies nutrient deprivation, resulting in longer-lived individuals (e.g. [76–78]). In addition, a suppressed IIS or removal of the germ-line produces life extending effects by activating the forkhead transcription factor (FOXO) which is conserved across *C. elegans* (*daf*-16), *D. melanogaster* (dFOXO) and mammals (FOXO3a) [79,80].

At the whole-body level [71], AMPK regulates metabolic energy balance by affecting feeding behaviour and circadian rhythms. When nutrient abundance is low, the elevated AMP to ATP ratio activates AMPK, with subsequent gain in health span and longevity in *D. melanogaster*. AMPK is a conserved modulator of lifespan in flies and mammals linking energy sensing to longevity, and is emerging as a major mechanism accounting for variation in longevity [71].

#### ii Lessons from studies with more ecological context

Hormone pathways have also been implicated in nutrient allocation shifts in non-model systems. Studies in flies and beetles have likewise suggested the IIS as a major pathway involved in resource distribution. An exonic indel polymorphism in the Insulin-like Receptor (*lnR*) gene was identified as a functional direct candidate target of natural selection in wild *D. melanogaster* [81,82]. In rhinoceros beetles, horn size is highly sensitive to nutrition and to perturbations in the IIS than are other body structures [83]. The precise details about how nutrients are mobilized toward competing traits have perhaps been best characterized in the wing dimorphic sand cricket, *Gryllus firmus* [84]. Juvenile hormone (JH) levels determine the morph, and trigger a whole host of processes leading to differential allocation of actual resource components toward flight capability versus reproduction. Flight capable morphs preferentially metabolize amino acids and convert a larger proportion of fatty acids to triglycerides while flightless morphs preferentially metabolize fatty acids and convert a larger proportion of amino acids to ovarian protein [65,84]. Adult crickets on low food diets allocate proportionally fewer resources toward flight capability [17,18], however, whether this diet-dependent shift is also mediated through JH has not yet been established. Juvenile hormone signalling is also involved in nutrition-based sex-specific mandible development via *doublesex* gene in the staghorn beetle [62].

These studies support the hypothesis that the evolution of allocation patterns ultimately results from the evolution of key endocrine pathways [66–68], potentially providing a simple theme in complex web of traits at various levels. Thus, while there is no denying acquisition and allocation of resources are highly complex processes, it is clear that hormone pathways serve as major mediators in many cases.

#### iii Understanding the underlying genetics of natural variation

Most of the above-described studies that identify key genes (except a few e.g. [81,82]) rely on evidence from large effect mutations or major perturbations and they have been very successful at identifying genes involved in the regulation of metabolism and resource allocation and of the effects of large alterations to individual genes. Our knowledge of the genetic basis of *natural* variation in metabolism and resource allocation is severely lacking in comparison, a predicament that is shared by the majority of complex traits [85–87]. The large effect genetic mutants identified via classical genetic techniques are typically not segregating in natural populations, which is not surprising given the central role of the pathways involved [88]. Additionally, despite the fact that several large effect mutations have been found to influence lifespan in *D. melanogaster* (see [3]), mapping studies and evolution experiments using natural populations have not independently identified these same genes as important contributors to natural genetic variation (e.g. [3,89–91]), with few exceptions [85,86]. These results are not due to a lack of genetic variation at these loci, given the populations used are derived from wild populations and typically have high heritabilities for most phenotypes, including gene expression levels of some of these same genes [92]. There are several possible explanations for this large disconnect regarding genes in these hormone pathways: 1) they do not contribute to natural genetic variation, 2) their effects are subtle and thus difficult to detect, 3) their effects stem from *trans* regulatory changes affecting gene expression [93]. Large effect mutant studies may represent the extreme tail of effect size distribution in nature [14], or, in the case of increase in longevity, different mechanisms altogether may induce altered nutrient signalling pathways in captive populations due to absence of stressors [86,87,94].

One of the strongest messages to emerge from modern quantitative genetics is that the genotype to phenotype map is more complex than some anticipated [95]. Within this complexity, our goals should be to find the main roads and general patterns. Newer mapping strategies, such as multi-parental populations, will help to assay multi-level traits on the same set of lines and leverage what is known about hormone pathways to reveal mechanistic bases of plastic resource allocation in natural organisms.

### b Integrating genetic and physiological mechanisms into evolutionary perspectives of resource allocation

As with the above evolutionary models, traditionally, questions surrounding proximate mechanisms have been considered separately from evolutionary questions, with a more recent movement toward integration across sub-disciplines. In particular, a major question surrounding hypothesized resource-based trade-offs is the degree to which the proximate mechanism underlying trade-offs stems from functional resource competition, or whether some other mechanism (e.g. hormone signalling), produces the relationship between traits. Here, we argue that these proximate mechanisms are not in conflict with the conceptual framework of the Y model.

#### i Challenge of a resource-based Y model

The Y model of resource allocation, as a framework to explain proximate mechanisms underlying life history trade-offs [12,96], has in recent years been criticized by some as inadequate, leading some to seek revision of life history theory (see exchanges in [97–101]). The challenge to a resource-cantered model is based on new empirical data showing that 1) abrogation of reproduction does not always extend lifespan, 2) some mutations that extend lifespan do not affect, or in fact, increase fecundity, and 3) male and female organisms of several species respond differently to interventions that increase lifespan. The most notable of these are studies in *C. elegans* [102,103] and *D. melanogaster* [104,105] in which gonad ablation failed to increase lifespan, while ablation of the germline only, doubled lifespan. Evidence suggests this effect is mediated largely by the insulin/IGF-1 system, which is thought to integrate molecular signals from the germ line and those from the somatic gonad to determine lifespan, rather than direct redistribution of resources. This hormonal signalling alternative has spurred a vigorous debate [99,100] whose reconciliation, in our view, depends on the eventual and successful integration of proximate mechanisms of trade-offs into evolutionary theory.

#### ii Is the new data really in conflict with the Y model?

We have reviewed above, case studies that directly or indirectly offer support for a resource model of life history evolution. Of particular note are studies demonstrating preferential amino acid metabolism and allocation of fatty acids to either flight or reproduction in winged vs wingless cricket morphs [84,106–110]. These works represent compelling evidence for differential resource allocation associated with the flight capability-reproduction trade-off. In addition, studies that quantified amino acid metabolism *in vivo* confirmed the predictions of the Y model for this trade-off [84,107]. Studies that fail to find the trade-off or find a positive relationship may not logically invalidate those that observe a negative correlation as multiple factors may be responsible. Further, the bulk of known mechanisms have been described in non-natural laboratory mutant organisms with limited or zero selection pressures experienced in the wild [86,87,94]. Although, the evidence for the connection between signalling and resource allocation is unclear, this absence of evidence should not be treated as evidence of absence.

#### iii Opportunities for integration of fields

Conceptual dichotomies where available empirical data do not sufficiently fit standing theoretical principles are not new to biology. These apparent conflicts have fuelled progress of the broad field and successfully led to the integration of once thought disparate fields – Mendelian, molecular, and quantitative genetics in the last century (see [111]). Instead of asking whether survival costs are best explained either by literal resource competition or by resource-free signalling, it may be useful to explore how the two integrate into the observed trade-off. This strategy can redirect attention to potential connections between nutrients and signals and factors that affect that connection. There is strong evidence that hormonal signalling is involved in nutrient sensing mechanisms implicated in aging [112], and that these mechanisms are at the base of appetite regulation and redistribution of nutrients [71]. It is thus possible to see how hormonal signalling may regulate optimal allocation and account for evolution of diverse resource allocation patterns. Thus, new data showing that signals regulate lifespan do not, presently, preclude the evolutionary role of resource constraints, especially in natural settings. It is completely fitting with evolutionary theory to expect organisms to use specific cues to indicate environmental conditions such as food availability. Thus, when results find that a single amino acid level can change how organisms allocate resources [113], an evolutionary interpretation is that that amino acid is what is cueing the organism about the environment, not that actual resource levels are not important to the response.

We do not yet know whether one or more proximate explanations govern a given trade-off. A possible scenario to emerge may negate the notion of a single proximate explanation since there may be unique mechanisms in different species and/or environmental settings. For example, in selected lines of *D. melanogaster*, offspring ovariole number increased in response to maternal starvation [114]; in wild-living *D. melanogaster* larval age and larval weight predicted survival in temperate and tropical regions, respectively [115]; and, in redback spider dietary restriction extended lifespan in mated, but imposed cost in unmated females [116]. These examples suggest multiple mechanistic possibilities defining a given phenotypic trade-off in different species or within species in different environments. Whether the trade-off is affected by diet, temperature, or behaviour, molecular signalling could lead to changes in how resources are allocated. It will benefit both fields if future studies take advantage of the ‘omic’ technologies to step up cross-field approaches in the search for mechanisms governing these traits in nature.

## 5 Future Directions

In this review, we have attempted to argue that a resource-based Y model is uniquely favoured to facilitate integration of evolutionary life history theory with proximate mechanisms underlying the near-ubiquitous trade-offs in life history traits. In doing so we have brought to the fore two key areas where significant progress is attainable, especially with the aid of ‘omic’ approaches: 1) performing studies in more ecologically-relevant contexts, and 2) increasing the level of integration between fields.

A major gap in our understanding of life history trade-offs in general, and the relationship between survival and reproduction in particular, is a general paucity of studies focusing on the underlying mechanisms in natural species, and lack of concordance between results of mutational studies in model species and those from studies of natural variation in the few cases where these have been undertaken. Here, we have attempted to show the wide variety of plastic resource allocation strategies in response to environmental fluctuations in availability that exist among natural populations and species. Understandably, many of the patterns so far uncovered have been demonstrated using laboratory studies with explicit diet manipulations (at most, three diet variations). We support this approach but, in addition, advocate for a broader, more realistic consideration of experimental diets. In this direction, studies taking ‘nutritional geometry’ approaches discussed earlier have the potential to provide a broader understanding of how organisms respond to changes in diet. In addition to considerations of mere nutritional content, experimental diets should attempt to mimic the natural diet of the organism as closely as possible, and the natural range of availability in the field in order to ensure that results most reflect evolutionarily relevant patterns that occur in nature.

Secondly, we have highlighted gaps in theoretical evolutionary models that address both optimal resource allocation patterns, and the evolution of phenotypic plasticity. To our knowledge, very few models specifically focus on evolution of phenotypically plastic resource allocation in response to variation in resource availability. We encourage the development of evolutionary models that integrate proximate mechanisms as a way to expand our understanding of the evolution of resource allocation strategies in multiple systems. In addition, testing the predictions of models predicting the evolution of different resource allocation strategies should be a major priority. Natural systems where patterns of availability differ among populations and species, can also inform these questions. Alternatively, experimental evolution approaches, where resource availability can be altered in a controlled way, and different types of variability across time can be induced, are potentially a powerful way to test these models. An experimental evolution approach could also allow for tracking change across the genotype to phenotype map in an integrative way, tracking changes in proximate mechanisms as evolution occurs.

Overall, viewing phenotypes within a framework of resource acquisition and allocation allows for a natural integration of physiology, genetics, and evolution. Studies that measure phenotypes at multiple levels (genomic, physiological, organismal levels) and in multiple resource environments provide a potentially productive path forward.

## Author’s contributions

EGK and EN conceived the idea and wrote the manuscript; AMP and EN compiled Table S1, EN, AMP and EGK provided editorial comments. All authors gave final approval for publication.

---

**Table S1:** Examples of the diversity of resource allocation strategies in some life history traits across the animal kingdom.

| Trade-off | Taxa | Increased allocation with low resource availability | Selected examples |
| --- | --- | --- | --- |
| **a) Trade-offs presented in the literature as two-trait cases** |  |  |  |
| Dispersal–reproduction | *Prokelisia marginata* | dispersal | [1] |
|  | *Gryllus rubens* | flightlessness | [2] |
|  | *Gryllus firmus* | flightlessness | [3] |
|  | *Gryllus firmus* | adults to dispersal;<br>juveniles to reproduction | [4,5] |
| Growth–reproduction* | *Daphnia magna* | growth | [6] |
| Current reproduction–future reproduction | *Synchaeta pectinata* | current reproduction<sup>^</sup> | [7] |
| Storage–reproduction | *Drosophila melanogaster* | storage | [8] |
| Survival–reproduction<sup>#</sup> | *Nothobranchius furzeri* | survival | [9] |
|  | *Trichoptera* spp. | to storage in short lived species; no change in long lived species | [10] |
|  | *Elephas maximus* | survival | [11] |
| | *Poecilia reticulata* | reproduction<sup>\$</sup> | [12] |
|  | Theraphosidae | survival | [13] |
|  | *Callosobruchus maculatus* | survival | [14] |
|  | *Asobara tabida* | survival | [15] |
|  | *Anastrepha ludens* | survival | [16] |
|  | *Notiophilus buguttatu* | survival | [17] |
|  | *D. melanogaster* | survival | [18–22] |
|  | *Rhabditophora* | reproduction | [23] |
|  | 10 rotifer species | most to survival | [24] |
|  | *Eupelmus vuilletti* | depends on nutrient (lipid vs sugar) | [25] |
|  | *Romalea microptera* | similar allocation to both traits | [26] |
|  | *Odocoileus virginianus* | survival | [27] |
|  | *Larus michahellis* | survival | [28] |
|  | *Saccopteryx bilineata* | survival | [29] |
|  | *Gerris* spp. | survival | [30] |
|  | *Diomedea exulans* | survival (with terminal reproductive investment) | [31] |
| Survival–body size | *Macaca mulatta*,<br>*Saimiri* sp. | M. mulatta to survival; Saimiri sp no effect | [32] |
| Starvation resistance–reproduction | *D. melanogaster* | starvation resistance | [18] |
| Egg mass–clutch size | *Microlophus delanonis* | to none | [33] |
| Total clutch mass–post-nesting condition | *Microlophus delanonis* | to none | [33] |
| Egg size–egg number | *Northobranchius furzeri* | egg size | [9] |
| Offspring quality–reproduction | examines 24 bird species on elevational gradient | quality in high elevation species | [34] |
| **b) Trade-offs presented in the literature as multi-trait cases** |  |  |  |
| Body condition–progeny quality<sup>‡</sup> | *Paroedura picta* | body condition | [35] |
| Dispersal–survival–reproduction–body mass | *Speyeria mormonica* | survival | [36] |
| Dispersal–lifespan–reproduction | *Colias eurytheme*,<br>*Speyeria mormonica* | dispersal, no effect on lifespan | [37] |
| Growth rate–development time–body size | *Scathophaga stercoraria* | faster growth rate and faster development time | [38] |
| Growth–reproduction– sprint speed | *Anolis sagrei* | survival | [39] |

<sup>‡</sup>Many of these refer to early reproductive effort; \*current reproduction; ^ovary size reduction, not absolute resource allocation; <sup>‡</sup>body length, head length vs clutch size, egg size; <sup>§</sup>Study measures somatic investment which may benefit both lifespan and fecundity.

## References

1. Sohal R, Weindruch R. 1996 Oxidative stress, caloric restriction, and aging. Sci. N. Y. NY

2. Browner WS, Kahn AJ, Ziv E, Reiner AP, Oshima J, Cawthon RM, Hsueh W-C, Cummings SR. 2004 The genetics of human longevity. Am. J. Med. 117, 851–860. (doi:10.1016/j.amjmed.2004.06.033)

3. Hughes KA, Reynolds RM. 2005 Evolutionary and mechanistic theories of aging. Annu. Rev. Entomol. 50, 421–445. (doi:10.1146/annurev.ento.50.071803.130409)

4. Kenyon CJ. 2005 The plasticity of aging: insights from long-lived mutants. Cell 120, 449–460. (doi:10.1016/j.cell.2005.02.002)

5. Partridge L, Alic N, Bjedov I, Piper MDW. 2010 Ageing in *Drosophila*: the role of the insulin/Igf and TOR signalling network. EXG 46, 1–6. (doi:10.1016/j.exger.2010.09.003)

6. Andersson M. 1986 Evolution of condition-dependent sex ornaments and mating preferences: sexual selection based on viability differences. Evolution 40, 804–816. (doi:10.2307/2408465)

7. Warren IA, Gotoh H, Dworkin IM, Emlen DJ, Lavine LC. 2013 A general mechanism for conditional expression of exaggerated sexually-selected traits. BioEssays News Rev. Mol. Cell. Dev. Biol. 35, 889–899. (doi:10.1002/bies.201300031)

8. Neel J. 1962 Diabetes mellitus: A ‘thrifty’ genotype rendered detrimental by ‘progress’? Am. J. Hum. Genet. 14, 353–362.

9. Wells JCK. 2009 Thrift: a guide to thrifty genes, thrifty phenotypes and thrifty norms. Int. J. Obes. 2005 33, 1331–1338. (doi:10.1038/ijo.2009.175)

10. Stearns SC. 1992 The evolution of life histories. Oxford University Press Oxford.

11. Roff DA, Mostowy S, Fairbairn DJ. 2002 The evolution of trade-offs: testing predictions on response to selection and environmental variation. Evol. Int. J. Org. Evol. 56, 84–95.

12. van Noordwijk AJ, de Jong G. 1986 Acquisition and allocation of resources: their influence on variation in life history tactics. Am. Nat. 128, 137–142.

13. Manolio TA et al. 2009 Finding the missing heritability of complex diseases. Nature 461, 747–753. (doi:10.1038/nature08494)

14. Rockman MV. 2012 The QTN program and the alleles that matter for evolution: all that’s gold does not glitter. Evolution 66, 1–17. (doi:10.1111/j.1558-5646.2011.01486.x)

15. Simmons F, Bradley T. 1997 An analysis of resource allocation in response to dietary yeast in *Drosophila melanogaster*. J. Insect Physiol. 43, 779–788.

16. Casas J, Pincebourde S, Mandon N, Vannier F, Poujol R, Giron D. 2005 Lifetime nutrient dynamics reveal simultaneous capital and income breeding in a parasitoid. Ecology 86, 545–554. (doi:10.1890/04–0812)

17. King EG, Roff DA, Fairbairn DJ. 2011 Trade-off acquisition and allocation in *Gryllus firmus*: a test of the Y model. J. Evol. Biol. 24, 256–264. (doi:10.1111/j.1420-9101.2010.02160.x)

18. King EG, Roff DA, Fairbairn DJ. 2011 The evolutionary genetics of acquisition and allocation in the wing dimorphic cricket, *Gryllus firmus*. Evolution 65, 2273–2285. (doi:10.1111/j.1558–5646.2011.01296.x)

19. Mockett RJ, Cooper TM, Orr WC, Sohal RS. 2006 Effects of caloric restriction are species-specific. Biogerontology 7, 157–160. (doi:10.1007/s10522–006-9004–3)

20. Kaitala A. 1991 Phenotypic plasticity in reproductive behavior of waterstriders: trade-offs between reproduction and longevity during food stress. Funct. Ecol. 5, 12–18.

21. Cooper TM, Mockett RJ, Sohal BH, Sohal RS, Orr WC. 2004 Effect of caloric restriction on life span of the housefly, *Musca domestica*. FASEB J. Off. Publ. Fed. Am. Soc. Exp. Biol. 18, 1591–1593. (doi:10.1096/fj.03–1464fje)

22. Weindruch R, Marriott BM, Conway J, Knapka JJ, Lane MA, Cutler RG, Roth GS, Ingram DK. 1995 Measures of body size and growth in rhesus and squirrel monkeys subjected to long-term dietary restriction. Am. J. Primatol. 35, 207–228.

23. Kirk KL et al. 2001 Dietary restriction and aging: comparative tests of evolutionary hypotheses. J. Gerontol. A. Biol. Sci. Med. Sci. 56, B123–9.

24. Stelzer C. 2001 Resource limitation and reproductive effort in a planktonic rotifer. Ecology 82, 2521–2533.

25. Calow P, Woollhead A. 1977 The relationship between ration, reproductive effort and age-specific mortality in the evolution of life-history strategies — some observations on freshwater triclads. J. Anim. Ecol. 46, 765–781.

26. Bashey F. 2006 Cross-generational environmental effects and the evolution of offspring size in the Trinidadian guppy *Poecilia reticulata*. Evolution 60, 348–361.

27. Harrison R G. 1980 Dispersal polymorphisms in insects. Annu. Rev. Ecol. Syst. 11, 95–118. (doi:10.1146/annurev.es.11.110180.000523)

28. Roff DA. 1986 The evolution of wing dimorphism in insects. Evolution 40, 1009–1020. (doi:10.2307/2408759)

29. Dingle H. 1996 Migration: the biology of life on the move: the biology of life on the move. California: Oxford University Press, USA.

30. Zera AJ, Denno RF. 1997 Physiology and ecology of dispersal polymorphism in insects. Annu. Rev. Entomol. 42, 207–230.

31. Denno RF, Douglas LW, Jacobs D. 1985 Crowding and host plant nutrition: environmental determinants of wing-form in Prokelisia marginata. Ecology 66, 1588–1596. (doi:10.2307/1938021)

32. Braendle C, Davis GK, Brisson JA, Stern DL. 2006 Wing dimorphism in aphids. Heredity 97, 192–199. (doi:10.1038/sj.hdy.6800863)

33. Zera AJ, Tiebel KC. 1988 Brachypterizing effect of group rearing, juvenile hormone III and methoprene in the wing-dimorphic cricket, *Gryllus rubens*. J. Insect Physiol. 34, 489–498. (doi:10.1016/0022–1910(88)90190–4)

34. Roff D. 1990 Antagonistic pleiotropy and the evolution of wing dimorphism in the sand cricket, *Gryllus firmus*. Heredity 65, 169–177.

35. Simpson SJ, Raubenheimer D. 1995 The geometric analysis of feeding and nutrition: a user’s guide. J. Insect Physiol. 41, 545–553. (doi:10.1016/0022-1910(95)00006-G)

36. Lee KP, Simpson SJ, Clissold FJ, Brooks R, Ballard JWO, Taylor PW, Soran N, Raubenheimer D. 2008 Lifespan and reproduction in *Drosophila*: new insights from nutritional geometry. Proc. Natl. Acad. Sci. U. S. A. 105, 2498–2503. (doi:10.1073/pnas.0710787105)

37. Behmer ST. 2009 Insect herbivore nutrient regulation. Annu. Rev. Entomol. 54, 165–187. (doi:10.1146/annurev.ento.54.110807.090537)

38. Clark RM, Zera AJ, Behmer ST. 2015 Nutritional physiology of life-history trade-offs: how food protein-carbohydrate content influences life-history traits in the wing-polymorphic cricket *Gryllus firmus*. J. Exp. Biol. 218, 298–308. (doi:10.1242/jeb.112888)

39. Cotter SC, Simpson SJ, Raubenheimer D, Wilson K. 2011 Macronutrient balance mediates trade-offs between immune function and life history traits. Funct. Ecol. 25, 186–198. (doi:10.1111/j.1365–2435.2010.01766.x)

40. Forster MJ, Morris P, Sohal RS. 2003 Genotype and age influence the effect of caloric intake on mortality in mice. FASEB J. 17, 690–692 (doi:10.1096/fj.02-0533fje)

41. Williams G. 1966 Natural selection, the costs of reproduction, and a refinement of Lack’s principle. Am. Nat. 100, 687.

42. Gadgil M, Bossert WH. 1970 Life historical consequences of natural selection. Am. Nat. 104, 1–24.

43. Charlesworth B. 1980 Evolution in age-structured populations. Cambridge, England: Cambridge University Press.

44. Perrin N, Sibly R. 1993 Dynamic models of energy allocation and investment. Annu. Rev. Ecol. Syst. 24, 379–410.

45. McNamara JM, Houston AI. 1996 State-dependent life histories. Nature 380, 215–221. (doi:10.1038/380215a0)

46. Roff DA. 2002 Life history evolution. Sinauer Associates, Inc. Sunderland, MA.

47. Via S, Lande R. 1985 Genotype-environment interaction and the evolution of phenotypic plasticity. Evolution 39, 505–522.

48. Kawecki T, Stearns S. 1993 The evolution of life histories in spatially heterogeneous environments: optimal reaction norms revisited. Evol. Ecol. 7, 155–174.

49. Dewitt TJ, Sih A, Wilson DS. 1998 Costs and limits of phenotypic plasticity. Trends Ecol. Evol. 13, 77–81.

50. Scheiner SM. 1998 The genetics of phenotypic plasticity. VII. Evolution in a spatially-structured environment. J. Evol. Biol. 11, 303–320. (doi:10.1046/j.1420-9101.1998.11030303.x)

51. De Jong G. 1999 Unpredictable selection in a structured population leads to local genetic differentiation in evolved reaction norms. J. Evol. Biol. 12, 839–851. (doi:10.1046/j.1420–9101.1999.00118.x)

52. de Jong G, Behera N. 2002 The influence of life-history differences on the evolution of reaction norms. Evol. Ecol. Res. 4, 1–25.

53. Ernande B, Dieckmann U. 2004 The evolution of phenotypic plasticity in spatially structured environments: implications of intraspecific competition, plasticity costs and environmental characteristics. J. Evol. Biol. 17, 613–628.

54. King EG, Roff DA. 2010 Modeling the evolution of phenotypic plasticity in resource allocation in wing-dimorphic insects. Am. Nat. 175, 702–716. (doi:10.1086/652434)

55. Fischer B, Dieckmann U, Taborsky B. 2010 When to store energy in a stochastic environment. Evolution 65, 1221–1232. (doi:10.1111/j.1558–5646.2010.01198.x)

56. Fischer B, Taborsky B, Dieckmann U. 2009 Unexpected patterns of plastic energy allocation in stochastic environments. Am. Nat. 173, E108–E120. (doi:10.1086/596536)

57. Welch AM, Semlitsch RD, Gerhardt HC. 1998 Call duration as an indicator of genetic quality in male gray tree frogs. Science 280, 1928–1930. (doi:10.1126/science.280.5371.1928)

58. Holzer B, Jacot A, Brinkhof MWG. 2003 Condition-dependent signaling affects male sexual attractiveness in field crickets, *Gryllus campestris*. Behav. Ecol. 14, 353–359. (doi:10.1093/beheco/14.3.353)

59. Hamilton WD, Zuk M. 1982 Heritable true fitness and bright birds: a role for parasites? Science 218, 384–387. (doi:10.1126/science.7123238)

60. Hunt J, Bussière LF, Jennions MD, Brooks R. 2004 What is genetic quality? Trends Ecol. Evol. 19, 329–333. (doi:10.1016/j.tree.2004.03.035)

61. Getty T. 2006 Sexually selected signals are not similar to sports handicaps. Trends Ecol. Evol. 21, 83–88. (doi:10.1016/j.tree.2005.10.016)

62. Gotoh H, Miyakawa H, Ishikawa A, Ishikawa Y, Sugime Y, Emlen DJ, Lavine LC, Miura T. 2014 Developmental link between sex and nutrition; doublesex regulates sex-specific mandible growth via juvenile hormone signaling in stag beetles. PLOS Genet. 10, e1004098. (doi:10.1371/journal.pgen.1004098)

63. Zahavi A. 1975 Mate selection — A selection for a handicap. J. Theor. Biol. 53, 205–214. (doi:10.1016/0022–5193(75)90111–3)

64. Mangel M, Munch SB. 2005 A life-history perspective on short-and long-term consequences of compensatory growth. Am. Nat. 166, E155–E176.

65. Zera AJ, Harshman LG. 2009 Laboratory selection studies of life-history physiology in insects. In Experimental evolution: Concepts, methods, and applications of selection experiments, pp. 236–281. University of California Press.

66. Flatt T, Tu M-P, Tatar M. 2005 Hormonal pleiotropy and the juvenile hormone regulation of *Drosophila* development and life history. BioEssays News Rev. Mol. Cell. Dev. Biol. 27, 999–1010. (doi:10.1002/bies.20290)

67. Tatar M, Bartke A, Antebi A. 2003 The endocrine regulation of aging by insulin-like signals. Sci. N. Y. NY 299, 1346–1351.

68. Zera AJ, Harshman LG, Williams TD. 2007 Evolutionary endocrinology: the developing synthesis between endocrinology and evolutionary genetics. Annu. Rev. Ecol. Evol. Syst. 38, 793–817. (doi:10.1146/annurev.ecolsys.38.091206.095615)

69. Teleman AA. 2010 Molecular mechanisms of metabolic regulation by insulin in *Drosophila*. Biochem. J. 425, 13–26. (doi:10.1042/BJ20091181)

70. Wells JC. 2010 The thrifty phenotype: an adaptation in growth or metabolism? Am. J. Hum. Biol. 23, 65–75. (doi:10.1002/ajhb.21100)

71. Hardie DG, Ross FA, Hawley SA. 2012 AMPK: a nutrient and energy sensor that maintains energy homeostasis. Nat. Rev. Mol. Cell Biol. 13, 251–262. (doi:10.1038/nrm3311)

72. French SS, Denise Dearing M, Demas GE. 2011 Leptin as a physiological mediator of energetic trade-offs in ecoimmunology: implications for disease. Integr. Comp. Biol. 51, 505–513. (doi:10.1093/icb/icr019)

73. Rajan A, Perrimon N. 2012 Drosophila cytokine unpaired 2 regulates physiological homeostasis by remotely controlling insulin secretion. Cell 151, 123–137. (doi:10.1016/j.cell.2012.08.019)

74. Mirth C, Truman JW, Riddiford LM. 2005 The role of the prothoracic gland in determining critical weight for metamorphosis in *Drosophila melanogaster*. Curr. Biol. 15, 1796–1807. (doi:10.1016/j.cub.2005.09.017)

75. Mendes CC, Mirth CK. 2016 Stage-specific plasticity in ovary size is regulated by insulin/insulin-like growth factor and ecdysone signaling in *Drosophila*. Genetics 202, 703–719. (doi:10.1534/genetics.115.179960)

76. Clancy DJ, Gems D, Harshman LG, Oldham S, Stocker H, Hafen E, Leevers SJ, Partridge L. 2001 Extension of life-span by loss of CHICO, a *Drosophila* insulin receptor substrate protein. Sci. N. Y. NY 292, 104–106. (doi:10.1126/science.1057991)

77. Tatar M, Kopelman A, Epstein D, Tu M, Yin C, Garofalo R. 2001 A mutant *Drosophila* insulin receptor homolog that extends life-span and impairs neuroendocrine function. Sci. N. Y. NY 292, 107–110.

78. Kenyon C, Chang J, Gensch E, Rudner A, Tabtiang R. 1993 A *C. elegans* mutant that lives twice as long as wild type. Nature 366, 461–464. (doi:10.1038/366461a0)

79. Puig O, Marr MT, Ruhf ML, Tjian R. 2003 Control of cell number by *Drosophila* FOXO: downstream and feedback regulation of the insulin receptor pathway. Genes Dev. 17, 2006. (doi:10.1101/gad.1098703)

80. Motta MC, Divecha N, Lemieux M, Kamel C, Chen D, Gu W, Bultsma Y, McBurney M, Guarente L. 2004 Mammalian SIRT1 represses forkhead transcription factors. Cell 116, 551–563.

81. Paaby AB, Blacket MJ, Hoffmann AA, Schmidt PS. 2010 Identification of a candidate adaptive polymorphism for *Drosophila* life history by parallel independent clines on two continents. Mol. Ecol. 19, 760–774. (doi:10.1111/j.1365–294X.2009.04508.x)

82. Paaby AB, Bergland AO, Behrman EL, Schmidt PS. 2014 A highly pleiotropic amino acid polymorphism in the *Drosophila* insulin receptor contributes to life-history adaptation. Evolution 68, 3395–3409. (doi:10.1111/evo.12546)

83. Emlen DJ, Warren IA, Johns A, Dworkin I, Lavine LC. 2012 A mechanism of extreme growth and reliable signaling in sexually selected ornaments and weapons. Science 337, 860–864. (doi:10.1126/science.1224286)

84. Zhao Z, Zera AJ. 2006 Biochemical basis of specialization for dispersal vs. reproduction in a wing-polymorphic cricket: Morph-specific metabolism of amino acids. J. Insect Physiol. 52, 646–658. (doi:10.1016/j.jinsphys.2006.03.003)

85. Reed LK et al. 2014 Systems genomics of metabolic phenotypes in wild-type *Drosophila melanogaster*. Genetics 197, 781–793. (doi:10.1534/genetics.114.163857)

86. Briga M, Verhulst S. 2015 What can long-lived mutants tell us about mechanisms causing aging and lifespan variation in natural environments? Exp. Gerontol. 71, 21–26. (doi:10.1016/j.exger.2015.09.002)

87. Savory FR, Benton TG, Varma V, Hope IA, Sait SM. 2014 Stressful environments can indirectly select for increased longevity. Ecol. Evol. 4, 1176–1185. (doi:10.1002/ece3.1013)

88. Van Voorhies WA, Curtsinger JW, Rose MR. 2006 Do longevity mutants always show trade-offs? Exp. Gerontol. 41, 1055–1058. (doi:10.1016/j.exger.2006.05.006)

89. Remolina SC, Chang PL, Leips J, Nuzhdin SV, Hughes KA. 2012 Genomic basis of aging and life-history evolution in *Drosophila melanogaster*. Evolution 66, 3390–3403. (doi:10.1111/j.1558–5646.2012.01710.x)

90. Magwire MM, Yamamoto A, Carbone MA, Roshina NV, Symonenko AV, Pasyukova EG, Morozova TV, Mackay TFC. 2010 Quantitative and molecular genetic analyses of mutations increasing *Drosophila* life span. PLOS Genet. 6, e1001037. (doi:10.1371/journal.pgen.1001037)

91. Burke MK, King EG, Shahrestani P, Rose MR, Long AD. 2014 Genome-wide association study of extreme longevity in *Drosophila melanogaster*. Genome Biol. Evol. 6, 1–11. (doi:10.1093/gbe/evt180)

92. Najarro MA, Hackett JL, Smith BR, Highfill CA, King EG, Long AD, Macdonald SJ. 2015 Identifying loci contributing to natural variation in xenobiotic resistance in *Drosophila*. PLOS Genet. 11, e1005663. (doi:10.1371/journal.pgen.1005663)

93. Pomp D, Allan MF, Wesolowski SR. 2004 Quantitative genomics: exploring the genetic architecture of complex trait predisposition. J. Anim. Sci. 82 E-Suppl, E300–312.

94. Holliday R, Rattan SIS. 2010 Longevity mutants do not establish any ‘new science’ of ageing. Biogerontology 11, 507–511. (doi:10.1007/s10522–010-9288–1)

95. Robinson MR, Wray NR, Visscher PM. 2014 Explaining additional genetic variation in complex traits. Trends Genet. TIG 30, 124–132. (doi:10.1016/j.tig.2014.02.003)

96. Boggs CL. 2009 Understanding insect life histories and senescence through a resource allocation lens. Funct. Ecol. 23, 27–37. (doi:10.1111/j.1365-2435.2009.01527.x)

97. Flatt T, Promislow DEL. 2007 Still pondering an age-old question. Science 318, 1255–1256. (doi:10.1126/science.1147491)

98. Flatt T. 2011 Survival costs of reproduction in *Drosophila*. Exp. Gerontol. 46, 369–375. (doi:10.1016/j.exger.2010.10.008)

99. Flatt T, Heyland A, Stearns SC. 2011 What mechanistic insights can or cannot contribute to life history evolution - an exchange between Stearns, Heyland, and Flatt. In Mechanisms of life history evolution: the genetics and physiology of life history traits and trade-offs. (eds T Flatt, A Heyland), New York: Oxford University Press.

100. Stearns SC. 2011 Does impressive progress on understanding mechanisms advance life history theory? In Mechanisms of life history evolution: the genetics and physiology of life history traits and trade-offs. (eds T Flatt, A Heyland), New York: Oxford University Press.

101. Edward DA, Chapman T. 2011 Mechanisms underlying reproductive trade-offs: costs of reproduction. In Mechanisms of life history evolution: the genetics and physiology of life history traits and trade-offs (eds T Flatt, A Heyland), New York: Oxford University Press.

102. Hsin H, Kenyon C. 1999 Signals from the reproductive system regulate the lifespan of *C. elegans*. Nature 399, 362–366. (doi:10.1038/20694)

103. Barnes AI, Partridge L. 2003 Costing reproduction. Anim. Behav. 66, 199–204. (doi:10.1006/anbe.2003.2122)

104. Barnes AI, Boone JM, Jacobson J, Partridge L, Chapman T. 2006 No extension of lifespan by ablation of germ line in *Drosophila*. Proc. R. Soc. B Biol. Sci. 273, 939–947. (doi:10.1098/rspb.2005.3388)

105. Flatt T, Min K-J, D’Alterio C, Villa-Cuesta E, Cumbers J, Lehmann R, Jones DL, Tatar M. 2008 *Drosophila*, germ-line modulation of insulin signaling and lifespan. Proc. Natl. Acad. Sci. U. S. A. 105, 6368–6373. (doi:10.1073/pnas.0709128105)

106. Mole S, Zera AJ. 1993 Differential allocation of resources underlies the dispersal-reproduction trade-off in the wing-dimorphic cricket, *Gryllus rubens*. Oecologia 93, 121–127.

107. Zera AJ, Zhao Z. 2006 Intermediary metabolism and life-history trade-offs: differential metabolism of amino acids underlies the dispersal-reproduction trade-off in a wing-polymorphic cricket. Am. Nat. 167, 889–900. (doi:10.1086/503578)

108. Zera AJ. 2005 Intermediary metabolism and life history trade-offs: lipid metabolism in lines of the wing-polymorphic cricket, *Gryllus firmus*, selected for flight capability vs. early age reproduction. Integr. Comp. Biol. 45, 511–524.

109. Zera AJ, Brink T. 2000 Nutrient absorption and utilization by wing and flight muscle morphs of the cricket *Gryllus firmus*: implications for the trade-off between flight capability and early reproduction. J. Insect Physiol. 46, 1207–1218.

110. Zera AJ, Larsen A. 2001 The metabolic basis of life history variation: genetic and phenotypic differences in lipid reserves among life history morphs of the wing-polymorphic cricket, *Gryllus firmus*. J. Insect Physiol. 47, 1147–1160.

111. Provine W B. 1971 The origins of theoretical population genetics. Chicago: University of Chicago Press.

112. López-Otín C, Blasco MA, Partridge L, Serrano M, Kroemer G. 2013 The hallmarks of aging. Cell 153, 1194–1217. (doi:10.1016/j.cell.2013.05.039)

113. Grandison RC, Piper MDW, Partridge L. 2009 Amino-acid imbalance explains extension of lifespan by dietary restriction in *Drosophila*. Nature 462, 1061–1064. (doi:10.1038/nature08619)

114. Wayne ML, Soundararajan U, Harshman LG. 2006 Environmental stress and reproduction in *Drosophila melanogaster*: starvation resistance, ovariole numbers and early age egg production. BMC Evol. Biol. 6, 57. (doi:10.1186/1471–2148-6–57)

115. Bochdanovits Z, de Jong G. 2003 Temperature dependent larval resource allocation shaping adult body size in *Drosophila melanogaster*. J. Evol. Biol. 16, 1159–1167. (doi:10.1046/j.1420–9101.2003.00621.x)

116. Stoltz JA, Hanna R, Andrade MCB. 2010 Longevity cost of remaining unmated under dietary restriction. Funct. Ecol. 24, 1270–1280. (doi:10.1111/j.1365-2435.2010.01729.x)

## References

1. Denno RF, Douglas LW, Jacobs D. 1985 Crowding and host plant nutrition: environmental determinants of wing-form in *Prokelisia marginata*. Ecology 66, 1588–1596. (doi:10.2307–1938021)

2. Zera AJ, Tiebel KC. 1988 Brachypterizing effect of group rearing, juvenile hormone III and methoprene in the wing-dimorphic cricket, *Gryllus rubens*. J. Insect Physiol. 34, 489–498. (doi:10.1016–0022-1910(88)90190–4)

3. Roff D. 1990 Antagonistic pleiotropy and the evolution of wing dimorphism in the sand cricket, *Gryllus firmus*. Heredity 65, 169–177.

4. King EG, Roff DA, Fairbairn DJ. 2011 Trade-off acquisition and allocation in *Gryllus firmus*: a test of the Y model. J. Evol. Biol. 24, 256–264. (doi:10.1111–j.1420–9101.2010.02160.x)

5. King EG, Roff DA, Fairbairn DJ. 2011 The evolutionary genetics of acquisition and allocation in the wing dimorphic cricket, *Gryllus firmus*. Evolution 65, 2273–2285. (doi:10.1111–j.1558-5646.2011.01296.x)

6. Yampolsky L, Ebert D. 1994 Variation and plasticity of biomass allocation in *Daphnia*. Funct. Ecol. 8, 435–440.

7. Stelzer C. 2001 Resource limitation and reproductive effort in a planktonic rotifer. Ecology 82, 2521–2533.

8. Djawdan M, Sugiyama TT, Schlaeger LK, Bradley TJ, Rose MR. 1996 Metabolic aspects of the trade-off between fecundity and longevity in *Drosophila melanogaster*. Physiol. Zool. 69, 1176–1195. (doi:10.1086–physzool.69.5.30164252)

9. Vrtílek M, Reichard M. 2014 Highly plastic resource allocation to growth and reproduction in females of an African annual fish. Ecol. Freshw. Fish 24, 616–628. (doi:10.1111–eff.12175)

10. Stevens DJ, Hansell MH, Monaghan P. 2000 Developmental trade-offs and life histories: strategic allocation of resources in caddis flies. Proc. R. Soc. B Biol. Sci. 267, 1511–1515.

11. Hayward AD, Mar KU, Lahdenperä M, Lummaa V. 2014 Early reproductive investment, senescence and lifetime reproductive success in female Asian elephants. J. Evol. Biol. 27, 772–783. (doi:10.1111–jeb.12350)

12. Bashey F. 2006 Cross-generational environmental effects and the evolution of offspring size in the Trinidadian guppy *Poecilia reticulata*. Evolution 60, 348–361.

13. Ibler B, Michalik P, Fischer K. 2013 Factors affecting lifespan in bird-eating spiders (Arachnida: Mygalomorphae, Theraphosidae) – A multi-species approach. Zool. Anz. 253, 126–136. (doi:10.1016–j.jcz.2013.09.004)

14. Messina F, Fry J. 2003 Environment-dependent reversal of a life history trade-off in the seed beetle *Callosobruchus maculatus*. J. Evol. Biol. 16, 501–509.

15. Ellers J, van Alphen JJM. 1997 Life history evolution in *Asobara tabida*: plasticity in allocation of fat reserves to survival and reproduction. J. Evol. Biol. 10, 771–785.

16. Carey JR, Harshman LG, Liedo P, Mller H-G, Wang J-L, Zhang Z. 2008 Longevity-fertility trade-offs in the tephritid fruit fly, *Anastrepha ludens*, across dietary-restriction gradients. Aging Cell 7, 470–477. (doi:10.1111–j.1474–9726.2008.00389.x)

17. Ernsting G, Isaaks JA. 1991 Accelerated ageing: a cost of reproduction in the carabid beetle *Notiophilus biguttatus* f. Funct. Ecol. 5, 299–303.

18. Chippindale A, Leroi AM, Kim S, Rose M. 1993 Phenotypic plasticity and selection in *Drosophila* life-history evolution. I. Nutrition and the cost of reproduction. J. Evol. Biol. 6, 171–193.

19. Good TP, Tatar M. 2001 Age-specific mortality and reproduction respond to adult dietary restriction in *Drosophila melanogaster*. J. Insect Physiol. 47, 1467–1473.

20. Sohal R, Weindruch R. 1996 Oxidative stress, caloric restriction, and aging. Sci. N. Y. NY

21. Partridge L, Piper MDW, Mair W. 2005 Dietary restriction in *Drosophila*. Mech. Ageing Dev. 126, 938–950. (doi:10.1016–j.mad.2005.03.023)

22. Lee KP, Simpson SJ, Clissold FJ, Brooks R, Ballard JWO, Taylor PW, Soran N, Raubenheimer D. 2008 Lifespan and reproduction in *Drosophila*: new insights from nutritional geometry. Proc. Natl. Acad. Sci. U. S. A. 105, 2498–2503. (doi:10.1073–pnas.0710787105)

23. Calow P, Woollhead A. 1977 The relationship between ration, reproductive effort and age-specific mortality in the evolution of life-history strategies — some observations on freshwater triclads. J. Anim. Ecol. 46, 765–781.

24. Kirk KL et al. 2001 Dietary restriction and aging: comparative tests of evolutionary hypotheses. J. Gerontol. A. Biol. Sci. Med. Sci. 56, B123–129.

25. Casas J, Pincebourde S, Mandon N, Vannier F, Poujol R, Giron D. 2005 Lifetime nutrient dynamics reveal simultaneous capital and income breeding in a parasitoid. Ecology 86, 545–554. (doi:10.1890–04–0812)

26. Heck MJ, Pehlivanovic M, Purcell JU, Hahn DA, Hatle JD. 2017 Life-extending dietary restriction reduces oxidative damage of proteins in grasshoppers but does not alter allocation of ingested nitrogen to somatic tissues. J. Gerontol. A. Biol. Sci. Med. Sci. 72, 616–623. (doi:10.1093–gerona– glw094)

27. Karns GR, Holland AM, Steury TD, Ditchkoff SS. 2014 Maternal life history of white-tailed deer: factors affecting fetal sex allocation, conception timing, and senescence. Evol. Ecol. Res. 16, 165–178.

28. Noguera JC, Lores M, Alonso-Álvarez C, Velando A. 2011 Thrifty development: early-life diet restriction reduces oxidative damage during later growth. Funct. Ecol. 25, 1144–1153. (doi:10.1111– j.1365–2435.2011.01856.x)

29. Greiner S, Nagy M, Mayer F, Knörnschild M, Hofer H, Voigt CC. 2014 Sex-biased senescence in a polygynous bat species. Ethology 120, 197–205. (doi:10.1111–eth.12193)

30. Kaitala A. 1991 Phenotypic plasticity in reproductive behavior of waterstriders: trade-offs between reproduction and longevity during food stress. Funct. Ecol. 5, 12–18.

31. Froy H, Phillips RA, Wood AG, Nussey DH, Lewis S. 2013 Age-related variation in reproductive traits in the wandering albatross: evidence for terminal improvement following senescence. Ecol. Lett. 16, 642–649. (doi:10.1111–ele.12092)

32. Weindruch R, Marriott BM, Conway J, Knapka JJ, Lane MA, Cutler RG, Roth GS, Ingram DK. 1995 Measures of body size and growth in rhesus and squirrel monkeys subjected to long-term dietary restriction. Am. J. Primatol. 35, 207–228.

33. Jordan M, Snell H. 2002 Life history trade-offs and phenotypic plasticity in the reproduction of Galapagos lava lizards (*Microlophus delanonis*). Oecologia 130, 44–52. (doi:10.1007–s004420100776)

34. Badyaev AV, Ghalambor CK. 2001 Evolution of life histories along elevational gradients: trade-off between parental care and fecundity. Ecology 82, 2948–2960. (doi:10.1890–0012-9658(2001)082[2948:EOLHAE]2.0.CO;2)

35. Kubička L, Kratochvíl L. 2009 First grow, then breed and finally get fat: hierarchical allocation to life-history traits in a lizard with invariant clutch size. Funct. Ecol. 23, 595–601. (doi:10.1111–j.1365-2435.2008.01518.x)

36. Niitepõld K, Boggs CL. 2015 Effects of increased flight on the energetics and life history of the butterfly *Speyeria mormonia*. PLOS ONE 10, e0140104. (doi:10.1371–journal.pone.0140104)

37. Niitepõld K, Perez A, Boggs CL. 2014 Aging, life span, and energetics under adult dietary restriction in lepidoptera. Physiol. Biochem. Zool. 87, 684–694. (doi:10.1086–677570)

38. Blanckenhorn WU. 1998 Adaptive phenotypic plasticity in growth, development, and body size in the yellow dung fly. Evolution 52, 1394–1407.

39. Cox RM, Calsbeek R. 2010 Severe costs of reproduction persist in *Anolis* lizards despite the evolution of a single-egg clutch. Evolution 64, 1321–1330. (doi:10.1111–j.1558–5646.2009.00906.x)

